# ACE2 expression in human dorsal root ganglion sensory neurons: implications for SARS-CoV-2 virus-induced neurological effects

**DOI:** 10.1101/2020.05.28.122374

**Authors:** Stephanie Shiers, Pradipta R. Ray, Andi Wangzhou, Claudio Esteves Tatsui, Larry Rhines, Yan Li, Megan L Uhelski, Patrick M. Dougherty, Theodore J Price

## Abstract

SARS-CoV-2 has created a global crisis. COVID-19, the disease caused by the virus, is characterized by pneumonia, respiratory distress and hypercoagulation and is often fatal^1^. An early sign of infection is loss of smell, taste and chemesthesis - loss of chemical sensation^2^. Other neurological effects of the disease have been described, but not explained^3,4^. We show that human dorsal root ganglion (DRG) neurons express the SARS-CoV-2 receptor^5,6^, ACE2. *ACE2* mRNA is expressed by a subset of nociceptors that express *MRGPRD* mRNA suggesting that SARS-CoV-2 may gain access to the nervous system through entry into neurons that form free-nerve endings at the outer-most layers of skin and luminal organs. Therefore, sensory neurons are a potential target for SARS-CoV-2 invasion of the nervous system.

## Introduction

The SARS-CoV-2 virus that causes COVID-19 enters cells via the angiotensin converting enzyme 2 (ACE2) receptor. The spike protein of the virus binds to ACE2^5,6^ and can be primed by a number of proteases (TMPRSS2^7^ and Furin^8^) that are sometimes co-expressed by ACE2 positive cells^9^. A set of genes that are involved in the response to SARS-CoV-2, SARS-CoV-2 and coronavirus-associated factors and receptors (SCARFs)^10^, have emerged allowing for assessment of virus effects on tissues through analysis of RNA sequencing or other high-throughput datasets.

Neurological symptoms are common in COVID-19 patients. Loss of smell (anosmia) is an early symptom which is now explained by non-neuronal ACE2 expression within the olfactory epithelium^11^. It is now appreciated that this anosmia is sometimes accompanied by loss of taste and loss of chemical sensation, chemesthesis^2^. This chemesthesis includes loss of capsaicin and menthol sensitivity^2^, sensations that are mediated by nociceptive sensory neurons^12^. Other neurological effects of SARS-CoV-2 associated with nociceptors have been described, including headache and nerve pain^1,4,13^. Molecular mechanisms underlying these effects are not known.

Bulk RNA sequencing datasets support the conclusion that *ACE2* is expressed in human DRG^14,15^, but cellular resolution is lacking. Single cell sequencing experiments from mouse DRG show weak *Ace2* expression in a subset of neurons that also express the *Mrgprd* and *Nppb* genes^16^. MrgprD is selectively expressed in a nociceptor population that forms free nerve endings in the skin^17^, cornea and luminal organs such as the colon^18^ as well as the meninges^19^. We tested the hypothesis that *ACE2* mRNA is expressed in a subset of human DRG neurons using tissues obtained from organ donors or vertebrectomy surgery based on RNAscope *in situ* hybridization. Our findings support the conclusion that approximately a quarter of human DRG neurons express *ACE2* and that most of these neurons are nociceptors that likely form free nerve endings in skin or other organs creating a potential entry point for the virus into the nervous system.

## Results

We conducted RNAscope for *ACE2* mRNA on human DRG to assess cellular expression of the SARS-CoV-2 receptor. We found that *ACE2* mRNA was localized in neurons, many of which expressed calcitonin gene-related peptide gene, *CALCA* (**Fig 1A-B**), and/or the P2X purinergic ion channel type 3 receptor gene, *P2RX3*. Both *CALCA* and *P2RX3* have been widely utilized to delineate nociceptor subpopulations in rodent DRG^12^. Expression of *ACE2* was consistent in lumbar DRG from a female organ donor (**Table 1**, demographic data for all DRGs analyzed) and in two thoracic DRGs taken from male patients undergoing vertebrectomy surgery (**Fig 1B**). Across samples, *ACE2* mRNA was present in 19.8 – 25.4% of all sensory neurons (**Fig 1C**), however, mRNA puncta for *ACE2* was sparse suggesting a low level of expression. Consistent with mouse single cell RNA sequencing data, *ACE2* neurons co-expressed *MRGPRD* and *NPPB mRNA* (**Fig 1D**). They also co-expressed Nav1.8 (*SCN10A* gene) mRNA (**Fig 1E**). *SCN10A* is a nociceptorspecific marker in rodent DRG at the protein and mRNA levels^20–22^; therefore, these findings are consistent with the notion that *ACE2* mRNA is mostly expressed by nociceptors in the human DRG. Summary statistics for the ACE2 population and *ACE2* mRNA-positive neuron cell sizes are shown in **Fig 1F** and **G**.

**Figure 1.**
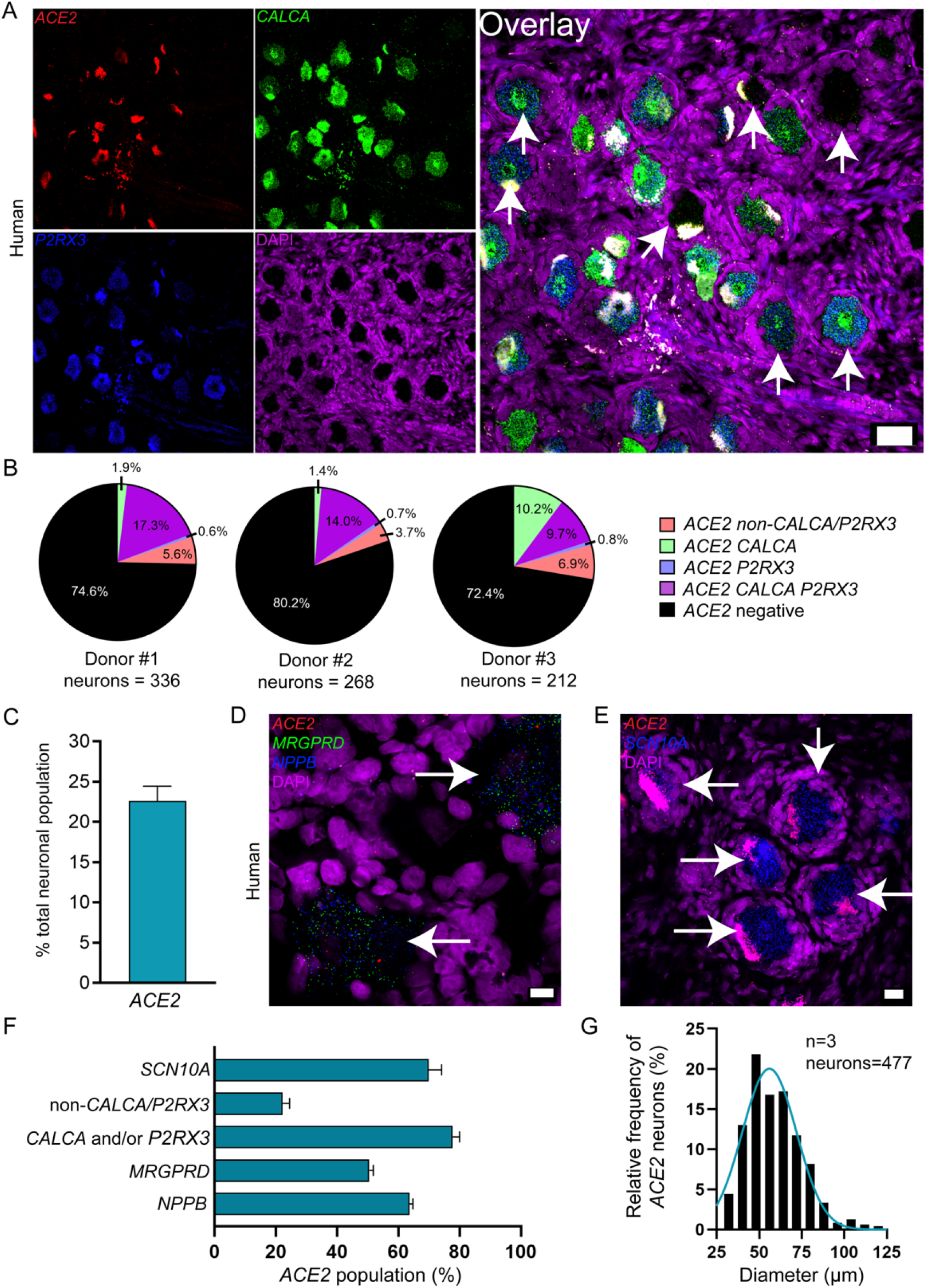
Distribution of ACE2 mRNA in human dorsal root ganglia. **A)** Representative 20x images of human DRG labeled with RNAscope in situ hybridization for CALCA (green), P2RX3 (blue), and ACE2 (red) mRNA and co-stained with DAPI (purple). Lipofuscin (globular structures) that autofluoresced in all 3 channels and appear white in the overlay image were not analyzed as this is background signal that is present in all human nervous tissue. **B)** Pie charts showing distribution of ACE2 neuronal subpopulations for each human donor DRG. ACE2 was found in neurons expressing solely CALCA (green), solely P2RX3 (blue) or both (purple), and a smaller population of neurons negative for both CALCA and P2RX3 (red, likely vascular or satellite glial but not immune cells, **Supplementary file 1**). **C)** ACE2 was expressed in 22.6% of all sensory neurons in human DRG. **D)** Representative 100x overlay image showing MRGPRD (green), NPPB (blue), ACE2 (red) and DAPI (purple) signal in human DRG. **E)** Representative 40x overlay image showing SCN10A (blue), ACE2 (red) and DAPI (purple) signal in human DRG. **F)** 69.9% of ACE2 neurons expressed SCN10A, 77.7% expressed CALCA and/or P2RX3, 22.2% were negative for CALCA and/or P2RX3, 50.4% expressed MRGPRD, and 63.6% expressed NPPB. Scale bars: 20x = 50μm, 40x = 20μm, 100x = 10μm. White arrows point toward ACE2-positive neurons.

**Table 1.** Human DRG tissue Information. Donor or patient information is given for all the samples that were used for RNAscope in situ hybridization shown in Fig 1.

| Donor # | DRG | Sex | Age | Race | Surgery/Cause of Death | Collection Site |
| --- | --- | --- | --- | --- | --- | --- |
| 1 | T4 | Male | 77 | White, non-Hispanic | Vertebrectomy | MD Anderson |
| 2 | L5 | Female | 33 | White, non-Hispanic | Opioid overdose | Southwest Transplant Alliance |
| 3 | T5 | Male | 56 | White, non-Hispanic | Vertebrectomy | MD Anderson |

Having established that the major receptor for SARS-CoV-2 is likely expressed by a specific subset of human nociceptors, we sought to use existing sequencing datasets to assess expression of other SCARFs^10^ in human DRG. We grouped these into receptors, proteases, replication factors, trafficking factors (shown only in **supplementary file 1** because they were all ubiquitous) and restriction factors (**Table 2** and **supplementary file 1**) as described in other surveys of RNA sequencing databases, none of which included DRG. While *ACE2* expression is low in these bulk RNA sequencing datasets, it was reliably detectable above 1 transcript per million (TPM) in certain thoracic DRG samples (**Supplementary file 1**). Some other potential receptors for SARS and MERS viruses were also found in human DRG and expressed in mouse DRG neurons, suggesting they may also be neuronally expressed in human DRG. Among proteases, the two main proteases for the SARS-CoV-2 spike protein priming, *TMPRSS2* and *FURIN*, were robustly expressed in human DRG and they were also expressed by mouse DRG neurons (**Table 2**). Most other SCARFs were also expressed in human DRG and, again, their expression in mouse DRG neurons suggests that most of them are likely to also be found in human DRG neurons. Many of these genes showed high variation between samples, an effect that may explain variable effects of SARS-CoV-2 on the nervous system in patients.

**Table 2.** Expression of SCARFs genes in human DRG from bulk RNA sequencing experiments. Genes are shown with mean +/- standard deviation (SD) expression in TPMs in 3 lumbar DRG samples and 5 thoracic DRG samples. Mouse DRG neuronal expression denotes detection in the neurofilament (NF), peptidergic (PEP), nonpeptidergic (NP) and tyrosine hydroxylase (TH) subsets of DRG neurons. Human DRG data is from^14,15^ and mouse single cell data is from^16^. ND: Not Detectable.

| Human gene | Lumbar DRG (mean +/- SD) TPM | Thoracic DRG (mean +/- SD) TPM | Neuronal expression in mouse DRG? | SCARF Classification |
| --- | --- | --- | --- | --- |
| *ACE2* | 0.09 +/- 0.02 | 0.94 +/- 0.88 | NF | Receptors |
| *BSG* | 304.82 +/- 82.3 | 239.42 +/- 79.23 | PEP, NP, NF, TH | Receptors |
| *ANPEP* | 14.56 +/- 7.27 | 13.73 +/- 4.26 | PEP, TH | Receptors |
| *CD209* | 0.57 +/- 0.43 | 1.60 +/- 1.28 | Multiple mouse orthologs | Receptors |
| *CLEC4G* | 0.79 +/- 0.34 | 0.22 +/- 0.45 | ND | Receptors |
| *CLEC4M* | 0.02 +/- 0.03 | 0.19 +/- 0.17 | No ortholog | Receptors |
| *DPP4* | 0.34 +/- 0.23 | 3.75 +/- 4.05 | ND | Receptors |
| *TMPRSS2* | 3.05 +/- 1.04 | 6.69 +/- 2.65 | NF, TH | Proteases |
| *TMPRSS11A* | ND | 0.06 +/- 0.13 | NF, TH | Proteases |
| *TMPRSS11B* | ND | 0.03 +/- 0.04 | No ortholog | Proteases |
| *FURIN* | 20.86 +/- 5.69 | 34.11 +/- 10.44 | PEP, NP, NF, TH | Proteases |
| *TMPRSS4* | 0.77 +/- 0.30 | 0.08 +/- 0.09 | NF | Proteases |
| *CTSL* | 154.53 +/- 21.71 | 105.19 +/- 13.36 | PEP, NP, NF, TH | Proteases |
| *CTSB* | 1116.17 +/- 132.45 | 697.50 +/- 96.96 | PEP, NP, NF, TH | Proteases |
| *TOP3B* | 6.46 +/- 0.88 | 18.93 +/- 5.45 | PEP, NP, NF, TH | Replication Factors |
| *MADP1* | 35.13 +/- 8.08 | 32.74 +/- 7.44 | PEP, NP, NF, TH | Replication Factors |
| *LY6E* | 446.65 +/- 234.34 | 218.17 +/- 161.15 | PEP, NP, NF, TH | Restriction Factors |
| *IFITM1* | 828.53 +/- 425.04 | 322.86 +/- 104.16 | PEP, NP, NF, TH | Restriction Factors |
| *IFITM2* | 880.05 +/- 340.01 | 2241.85 +/- 91.24 | PEP, NP, NF, TH | Restriction Factors |
| *IFITM3* | 2339.37 +/- 943.43 | 744.00 +/- 224.85 | PEP, NP, NF, TH | Restriction Factors |
| *CALCA* | 1634.46 +/- 665.44 | 524.86 +/- 111.60 | PEP, NP, NF, TH | Neuronal Markers |
| *P2RX3* | 44.65 +/- 42.69 | 144.54 +/- 47.60 | PEP, NP, NF, TH | Neuronal Markers |
| *NPPB* | 41.20 +/- 16.37 | 21.29 +/- 8.99 | NP, PEP | Neuronal Markers |
| *MRGPRD* | 3.01 +/- 4.62 | 10.47 +/- 7.64 | PEP, NP, NF, TH | Neuronal Markers |

## Discussion

Our work highlights neuronal expression of *ACE2* in a select subset of nociceptors that express *CALCA*, *P2RX3, MRGPRD, NPPB and SCN10A*. While we cannot state with certainty the anatomical projections of these neurons because tracing studies cannot be done in humans, the neurochemical signature of these neurons is consistent with nociceptors that form free nerve endings in the skin^17^, luminal organs^18^ and meninges^19^. Therefore, one potential consequence of this ACE2 expression could be infection of nociceptors through the nasal passages, cornea, or upper or lower airway. To this end, we noted higher expression of *ACE2* in thoracic DRGs, and these DRGs contain nociceptors that innervate the lungs^23,24^, a major site for proliferation of the SARS-CoV-2 virus^1^.

It has recently been recognized that an early symptom of SARS-CoV-2 infection is loss of smell and taste^2,13,25^. This has now been explained by SARS-CoV-2 infection of non-neuronal cells in the olfactory epithelium^25^. Another interesting early symptom of SARS-CoV-2 is chemesthesis^2^. This symptom is less frequent than loss of smell and cannot be explained via the same mechanisms as anosmia because the sensing of noxious chemicals in the oral cavity is mediated by nociceptors. Our data suggests that a possible explanation of chemesthesis in COVID-19 is silencing of nerve endings by viral infection. If pulmonary afferents are also infected by SARS-CoV-2, as our data would predict, this could have important consequences for disease severity. Previous studies have shown that ablation of airway nociceptors increases the severity of respiratory disease due to a loss of the trophic action of CGRP within the airway^26^. Widespread silencing of nociceptors innervating the oral and nasal cavity as well as the upper and lower airway may have important impacts on COVID-19 disease severity.

There are many currently unexplained neurological features in COVID-19 patients^4,13^. It is mysterious how the virus enters the nervous system as most neurons do not express ACE2^10,27^. Our work presents a plausible entry point – through a select subset of ACE2 expressing nociceptors that then form synapses with spinal and brainstem CNS neurons. While our findings can be taken as putative evidence for this idea, ACE2 expression is very low in these neurons, and we have not tested whether ACE2 protein can be detected in free nerve endings in appropriate organs, and whether such expression can be harnessed by the virus to gain entry. More work will be needed to make these determinations, but our results lay a foundation for understanding the symptoms, pathology and long-term outcomes of COVID-19 as they relate to the sensory nervous system.

## Materials and Methods

### Tissue preparation

All human tissue procurement procedures were approved by the Institutional Review Boards at the University of Texas at Dallas and University of Texas MD Anderson Cancer Center. Human dorsal root ganglion (at levels T4, T5, and L5) were collected, frozen on dry ice (L5) or liquid nitrogen (T4 and T5) and stored in a −80°C freezer. Donor and/or patient information is provided in **Table 1**. The human DRGs were gradually embedded with OCT in a cryomold by adding small volumes of OCT over dry ice to avoid thawing. All tissues were cryostat sectioned at 20 μm onto SuperFrost Plus charged slides. Sections were only briefly thawed in order to adhere to the slide but were immediately returned to the −20°C cryostat chamber until completion of sectioning. The slides were then immediately utilized for histology.

### RNAscope in situ hybridization

RNAscope *in situ* hybridization multiplex version 1 was performed as instructed by Advanced Cell Diagnostics (ACD). Slides were removed from the cryostat and immediately transferred to cold (4°C) 10% formalin for 15 minutes. The tissues were then dehydrated in 50% ethanol (5 min), 70% ethanol (5 min) and 100% ethanol (10 min) at room temperature. The slides were air dried briefly and then boundaries were drawn around each section using a hydrophobic pen (ImmEdge PAP pen; Vector Labs). When hydrophobic boundaries had dried, protease IV reagent was added to each section until fully covered and incubated for 5 minutes at room temperature. The protease IV incubation period was optimized as recommended by ACD for the specific lot of Protease IV reagent. Slides were washed briefly in 1X phosphate buffered saline (PBS, pH 7.4) at room temperature. Each slide was then placed in a prewarmed humidity control tray (ACD) containing dampened filter paper and a mixture of Channel 1, Channel 2, and Channel 3 probes (50:1:1 dilution, as directed by ACD due to stock concentrations) was pipetted onto each section until fully submerged. This was performed one slide at a time to avoid liquid evaporation and section drying. The humidity control tray was placed in a HybEZ oven (ACD) for 2 hours at 40°C. A table of all probes used is shown in **Table 3**. Following probe incubation, the slides were washed two times in 1X RNAscope wash buffer and returned to the oven for 30 minutes after submersion in AMP-1 reagent. Washes and amplification were repeated using AMP-2, AMP-3 and AMP-4 reagents with a 15-min, 30-min, and 15-min incubation period, respectively. AMP-4 ALT C (Channel 1 = Atto 550, Channel 2 = Atto 647, Channel 3 = Alexa 488) was used for all experiments. Slides were then washed two times in 0.1M phosphate buffer (PB, pH7.4). Human slides were incubated in DAPI (ACD) for 1 min before being washed, air dried, and cover-slipped with Prolong Gold Antifade mounting medium.

**Table 3.** Summary table of Advanced Cell Diagnostics (ACD) RNAscope probes.

| **mRNA** | **Gene name** | **ACD Probe Cat No.** |
| --- | --- | --- |
| *CALCA* | Calcitonin gene-related peptide | 605551-C2 |
| *P2RX3* | Purinergic Receptor P2X3 | 406301-C3 |
| *SCN10A* | Sodium Voltage-Gated Channel Alpha Subunit 10; Nav1.8 | 406291-C2 |
| *MRGPRD* | MAS Related GPR Family Member D | 524871-C3 |
| *NPPB* | Natriuretic Peptide B | 448511-C2 |
| *ACE2* | Angiotensin I Converting Enzyme 2 | 848151-C1 |

### Tissue Quality Check

All tissues were checked for RNA quality by using a positive control probe cocktail (ACD) which contains probes for high, medium and low-expressing mRNAs that are present in all cells (ubiquitin C > Peptidyl-prolyl cis-trans isomerase B > DNA-directed RNA polymerase II subunit RPB1). All tissues showed signal for all 3 positive control probes (**Supplementary Fig 1**). A negative control probe against the bacterial DapB gene (ACD) was used to check for non-specific/background label.

### Image Analysis

DRG sections were imaged on an Olympus FV3000 confocal microscope at 20X or 40X magnification. For the *CALCA/P2RX3/ACE2* and *SCN10A/ACE2* experiments, 3-4 20X images were acquired of each human DRG section, and 3-4 sections were imaged per human donor. For the *MRGPRD/NPPB/ACE2* experiments, 8 40x images were acquired of each human DRG section, and 3-4 sections were imaged per donor. The sections imaged were chosen at random, but preference was given to sections that did not have any sectioning artifact, or sections that encompassed the entire DRG bulb. The acquisition parameters were set based on guidelines for the FV3000 provided by Olympus. In particular, the gain was kept at the default setting 1, HV ≤ 600, offset = 4 (based on HI-LO settings; doesn’t change between experiments), and laser power ≤ 10% (but generally the laser power was ≤ 5% for our experiments). The raw image files were brightened and contrasted in Olympus CellSens software (v1.18), and then analyzed manually one cell at a time for expression of each gene target. Cell diameters were measured using the polyline tool. Total neuron counts for human samples were acquired by counting all of the probe-labeled neurons and all neurons that were clearly outlined by DAPI (satellite cell) signal and contained lipofuscin in the overlay image.

Large globular structures and/or signal that auto-fluoresced in all 3 channels (488, 550, and 647; appears white in the overlay images) was considered to be background lipofuscin and was not analyzed. Aside from adjusting brightness/contrast, we performed no digital image processing to subtract background. We attempted to optimize automated imaging analysis tools for our purposes, but these tools were designed to work with fresh, low background rodent tissues, not human samples taken from older organ donors. As such, we chose to implement a manual approach in our imaging analysis in which we used our own judgement of the negative/positive controls and target images to assess mRNA label. Images were not analyzed in a blinded fashion.

### RNA Sequencing Sources

We used previously published RNA sequencing datasets to generate **Table 2** and **Supplementary file 1**. Human DRG sequencing data was described previously^14,15^ and mouse single cell sequencing data from DRG was described previously^16,28^.

### Data Analysis

Graphs were generated using GraphPad Prism version 7.01 (GraphPad Software, Inc. San Diego, CA USA). Given that the percentage of *ACE2*-expressing neurons was assessed in each experiment, we averaged these numbers for each donor to generate the final data values. A total of 2,224 neurons were analyzed between all three donors and all three experiments. A relative frequency distribution histogram with a fitted Gaussian distribution curve was generated using the diameters of all *ACE2*-positive neurons detected in all experiments.

---

**Supplemental Figure 1.**
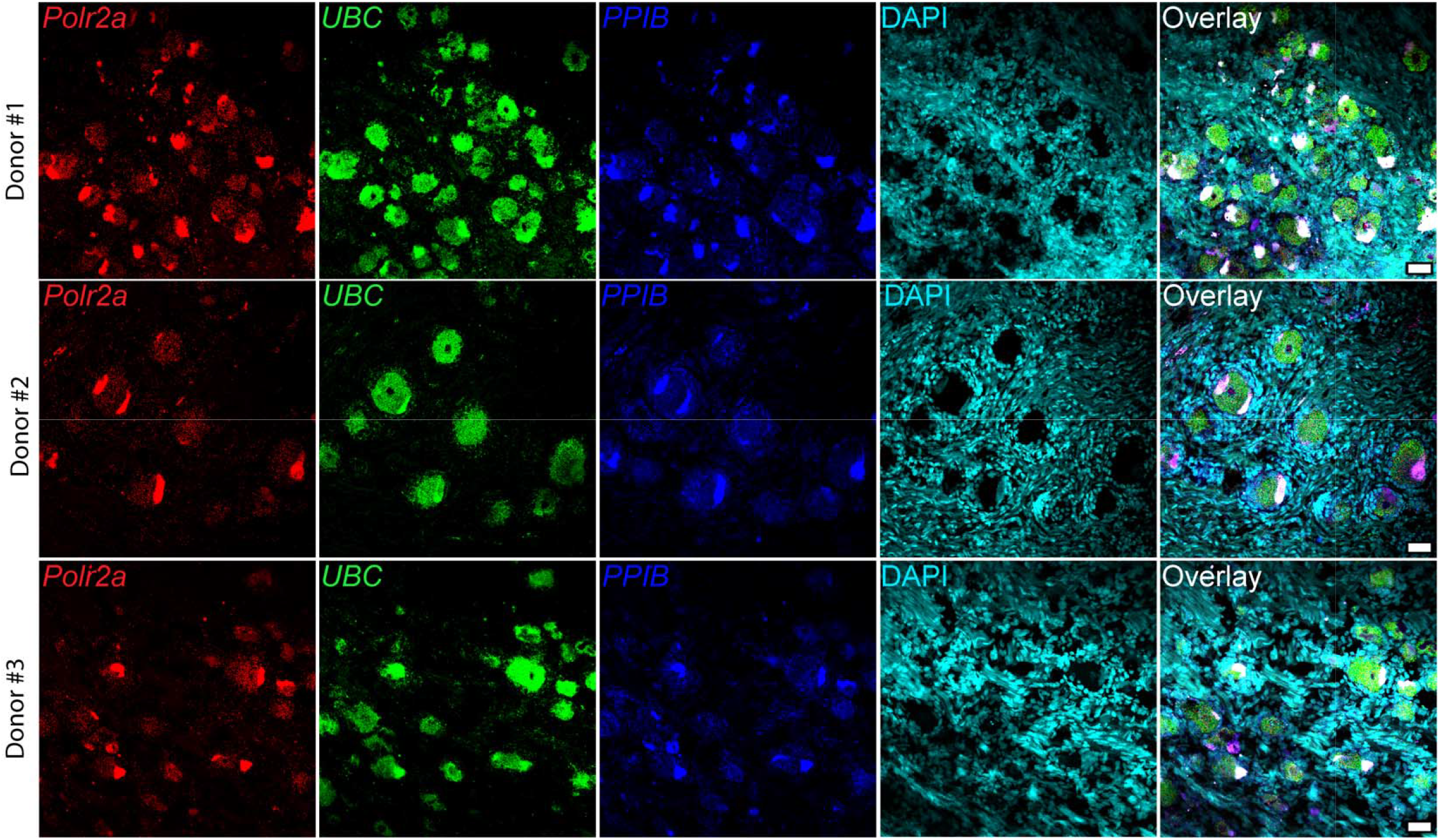
mRNA quality check for each human donor DRG. All tissues were checked for mRNA quality by using a positive control probe cocktail (ACD) which contains probes for high, medium and low-expressing mRNAs that are present in all cells (ubiquitin C > Peptidyl-prolyl cis-trans isomerase B > DNA-directed RNA polymerase II subunit RPB1). All DRGs showed signal for all 3 positive control probes indicating that the mRNA quality is excellent and that low-expressing transcripts such as *ACE2* should be able to be detected. 20x scale bar = 50 μm.

